# Temporal and Clonal Progression in a Pediatric Ependymoma Patient Through Multiple Treatments

**DOI:** 10.1101/115923

**Authors:** Christopher A. Miller, Sonika Dahiya, Tiandao Li, Robert S. Fulton, Matthew D. Smyth, Gavin P. Dunn, Joshua B. Rubin, Elaine R. Mardis

**Author notes:** Authors for correspondence: Elaine R. Mardis, The Institute for Genomic Medicine at Nationwide Children’s Hospital, The Ohio State University College of Medicine, 700 Children’s Drive, Columbus, OH 43205, Joshua B. Rubin, Department of Pediatrics, Washington University School of Medicine, Campus Box 8116 660 S. Euclid Ave, St. Louis, MO 63110.

## Abstract

**Background:** Multiple recurrences after complete resection and irradiation of supratentorial ependymoma are common and frequently result in patient death. However, the molecular basis for treatment resistance, the impact that radiation and other adjuvant therapies have in promoting recurrence, and the use of this information to rationally design effective approaches to treat recurrent ependymoma are unknown. Due to the rarity of these tumors and the even less likely banking of multiple recurrent samples from the same patient, we initiated a study to characterize the evolution of a single patient’s ependymoma in response to therapy.

**Methods and Findings:** A combination of high depth, whole genome and exome-based DNA sequencing of germline and tumor specimens, RNA sequencing of tumor specimens, and advanced computational analyses were employed to reconstruct the natural history of a supratentorial ependymoma case in which there were four local recurrences. The findings reveal the extent to which treatment with radiation and chemotherapies resulted in the diversification of the tumor subclonal architecture and shaped the neo-antigen landscape, and provide new insights into possible molecular mechanisms of oncogenesis, treatment response and recurrence.

**Conclusions:** Although the recurrent tumors we studied were clearly shaped by therapy, the founding clone was never eradicated by any treatment. We conclude that DNA and RNA sequencing may provide critical prognostic indicators to identify ependymoma patients that should be observed, rather than irradiated, post gross total resection.

## Background

Ependymomas are a heterogeneous group of primary central nervous system tumors with multiple histological, brain region, age and molecular features distinguishing between different prognostic groups [1][2][3]]. Overall, 10 year survival for children and adults with ependymomas is approximately 80%, with specific subsets of patients exhibiting survival of less than 30% at 10 years. Based on standard histological features, ependymal neoplasms can be diagnosed as World Health Organization (WHO) grade I, II or III tumors. However, in contrast to other brain tumors, histological grading has proven to be a weak prognostic indicator of outcome for ependymomas treated with standard-of-care complete surgical resection, with or without post-operative radiation therapy as indicated by the WHO grade [4]. Thus, there has been an increasing emphasis on the identification of prognostic molecular features that might provide more precise stratification for the risk of recurrence. For instance, young males harboring posterior fossa ependymomas found to exhibit a hypermethylator phenotype demonstrate poor survival [5]. Because the clinically distinguishing features of ependymoma include relatively common occurrence of multiple and late relapses, it is clear that improving outcomes and decreasing recurrence events will require a more complete understanding of tumor evolution after treatment.

To date, there is a paucity of information regarding the genomic changes in ependymomas that recur serially through multiple treatment regimens. This fact is largely due to the rarity of the disease and failure to bank and analyze recurrent samples. In order to determine the temporal genomic changes that occurred in one patient’s ependymoma disease as it recurred after several different therapeutic modalities, we characterized the genomic landscape of serial resections with high depth next generation whole genome and exome sequencing. These data provided an evaluation of putative driver mutations, mutational signatures resulting from therapy, mechanisms for therapy response and resistance, and shifts in the neoantigen profile from the initial disease presentation through four recurrences.

## Methods

### Case history

The initial diagnosis was made in a 16 year-old right-handed female who presented to the St Louis Children’s Hospital Emergency Department with a three-day history of headache and vomiting. MRI scan revealed a 6 cm × 4 cm enhancing mass in the right frontotemporal region (**Fig 1A initial diagnosis**). The patient underwent a gross total resection via a right frontotemporal craniotomy. Pathological evaluation was significant for a hypercellular glial tumor with prominent pseudo-rosettes, increased mitoses, vascular proliferation and necrosis, and perinuclear dot-like expression of epithelial membrane antigen (EMA, **Fig 1 B, C**) along-with diffuse glial fibrillary acidic protein immunoreactivity. A diagnosis of anaplastic ependymoma (WHO grade III) was made. Evaluations for central nervous system (CNS) dissemination were negative. The patient received 59.4 Gy of fractionated photon irradiation to the tumor bed plus a 1 cm margin, which is standard for supratentorial ependymoma. Forty-four months after the initial diagnosis, the patient suffered a seizure and an MRI revealed a 13 × 15 × 16 mm nodular recurrence in the right frontal lobe along the posterior margin of the initial resection cavity (**Fig 1A first recurrence**). MRI of spine and cerebrospinal fluid cytology were negative. The patient underwent complete resection of the recurrent tumor, which exhibited similar histology to the initial tumor. The resection cavity and margin were re-irradiated with an additional 59.4 Gy of fractionated photon irradiation and the patient received 10 months of standard dose temozolomide treatment.

**Fig 1.**
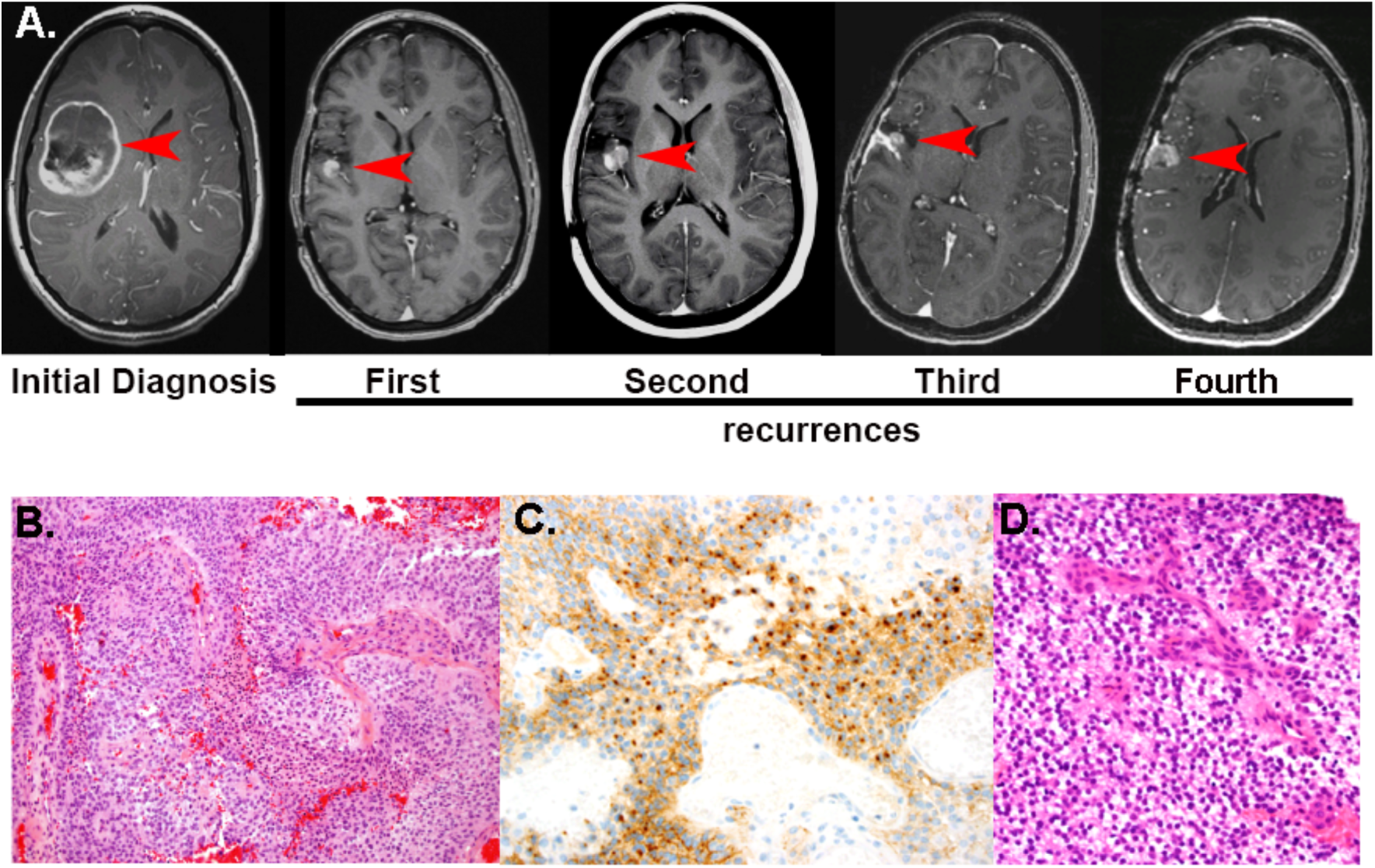
Radiographic and Pathological Evaluation of Initial and Recurrent Ependymoma. (**A**) Serial MRIs over a 10 year period demonstrating a heterogeneously enhancing mass in the right frontotemporal region at the time of initial diagnosis and four enhancing recurrent lesions adjacent to the initial resection cavity. (**B**) Hematoxylin and eosin stain of formalin fixed paraffin embedded primary resection material revealed a densely cellular tumor with increased mitotic activity, necrosis, and microvascular proliferation. (**C**) Immunostain for epithelial membrane antigen (EMA) shows multifocal perinuclear dot-like positivity, which is characteristic of ependymal differentiation. (**D**). Hematoxylin and eosin stain of each recurrent tumor revealed persistence of the ependymal phenotype. Pictured is the third recurrence. All the photomicrographs are taken at 40X magnification.

A surveillance scan 17 months after the second resection demonstrated a 1 cm enhancing nodule in the temporal surface of the right sylvian fissure near the resection cavity, consistent with recurrence (**Fig 1A second recurrence**). Following a third complete resection, histopathology was again consistent with anaplastic ependymoma and analysis for dissemination was negative. The patient was enrolled on CERN- 0801 at Children’s Memorial Hospital in Chicago and received combined Avastin and Lapatinib. Lapatinib was discontinued four months later due to toxicity and Avastin was continued for an additional eight months for a total of one year of treatment every two weeks. Six months later, an MRI revealed a new right peri-sylvian lesion and right thalamic enhancing nodule (**Fig 1A third recurrence**). Complete resection of the peri-sylvian lesion was performed and pathology again indicated anaplastic ependymoma (**Fig 1D**) with no evidence of dissemination. Avastin was restarted and continued for 20 months until new evidence from serial MRI indicated progression in a peri-sylvian lesion that had remained following the most recent surgery (**Fig 1A fourth recurrence**). This lesion also was completely resected and diagnosed as anaplastic ependymoma.

### DNA sequencing

DNA was isolated from fresh frozen sections of each tumor resection using the Qiagen Dual Prep, and evaluated for quality and concentration using established methods. DNA was isolated from PBMC after Ficoll-based isolation to provide a normal comparator, and evaluated for quality and concentration. Using 500 ng input for all five tumors and the blood normal DNA, we generated two indexed whole genome sequencing libraries by standard methods (Kapa Biosystems) for each sample. One library per sample was processed through exome hybrid capture using the IDT EzExome reagent (Integrated DNA Technologies, Coralville IA), quantitated and amplified post-capture using the manufacturer’s protocol. Each of the corresponding WGS libraries was amplified by PCR, quantitated and diluted as appropriate for Illumina sequencing. The final libraries for each sample (WGS + exome) were pooled to produce combined tumor and normal WGS and exome sequencing data in a specific proportion, yielding ~10-fold WGS and ~1000-fold exome coverage (**S1 Table**). The resulting library pools were loaded onto the HiSeqX platform and sequenced using 150 bp paired end reads.

## Somatic variant analysis

After index-based binning of the reads into WGS- and exome-derived tumor and normal data, sequence data were aligned to reference sequence build GRCh37-lite-build37 using BWA mem[ref] version 0.7.10 (params: -t 8::), then merged and deduplicated using picard version 1.113, (https://broadinstitute.github.io/picard/). Somatic variants were called from the combined data using our Genome Modeling System (GMS) [6] as follows:

SNVs were detected using the union of four callers: 1) samtools[7] version r982 (params: mpileup -BuDs) intersected with Somatic Sniper [8] version 1.0.4 (params: -F vcf -G –L -q 1 -Q 15) and processed through false-positive filter v1 (params: --bam-readcount- version 0.4 --bam-readcount-min-base-quality 15 --minmapping-quality 40 --min-somatic-score 40), 2) VarScan[9] version 2.3.6 filtered by varscan-high-confidence filter version v1 and processed through false-positive filter v1 (params: --bam-readcount-version 0.4 --bam-readcount-min-base-quality 15), 3) Strelka [10] version 1.0.11 (params: isSkipDepthFilters = 0), and 4) mutect [11] v1.1.4.

Indels were detected using the union of 4 callers: 1) GATK [12] somatic-indel version 5336) pindel version 0.5 filtered with pindel [13] somatic calls and VAF filters (params: --variant-freq-cutoff=0.08), and pindel read support, 3) VarScan version 2.3.6 filtered by varscan-high-confidence- indel version v1 and 4) Strelka version 1.0.11 (params: isSkipDepthFilters = 0).

SNVs and Indels were further filtered by removing artifacts found in a panel of 905 normal exomes, removing sites that exceeded 0.1% frequency in the 1000 genomes or NHLBI exome sequencing projects, and then using a Bayesian classifier (https://github.com/genome/genome/blob/master/lib/perl/Genome/Model/Tools/Validation/IdentifyOutliers.pm) and retaining variants classified as somatic with a binomial log-likelihood of at least 10.

For protein-coding mutation counts described in the results below, a variant was considered to be present in a sample if it appeared with at least 3 variant supporting-reads and a VAF of > 2.5%. As some sites had low or variable coverage, a variant was only considered to be completely cleared if it did not appear in any subsequent samples.

Copy number aberrations were detected using bam-window (window-size 10000) and copy-cat version 1.6.11 (params: --per-read-length –per-library) (https://github.com/chrisamiller/copyCat). Uneven sequence coverage of the normal sample required us to run copyCat in tumor-only mode, followed by manual review to differentiate somatic from germline copy number events.

Putative structural variants were detected using the union of Breakdancer 1.4.5 ^[14]^filtered by novo-realign and tigra-sv, and squaredancer 0.1 (https://github.com/genome/genome/blob/master/lib/perl/Genome/Model/Tools/Sv/SquareDancer.pl).

### RNA sequencing

Total RNA was concurrently isolated from each fresh frozen tumor resection (Qiagen Dual Prep), and evaluated for quality and concentration using the Agilent Tapestation. RNAseq libraries were constructed using the TruSeq Stranded RNAseq library kit (Illumina, Inc., San Diego CA) according to the manufacturer’s protocol, quantitated and diluted for sequencing. Using the HiSeq 2500, we produced sequencing data from each RNAseq library in a single flow cell lane by paired end 100 bp reads, yielding between 96 and 655 million reads per sample. The fourth recurrence sample was subjected to a capture step before sequencing, using the IDT EzExome reagent. This sample yielded 856 million reads, with a much higher coding-region percentage. (**S1 Table**).

### RNAseq analysis

The resulting read data were aligned to the human reference with Tophat v2.0.8 (denovo mode, params: -library-type fr-firststrand --bowtie-version=2.1.0). Expression levels were calculated with Cufflinks v2.1.1 (params: --max-bundle-length 10000000 --max-bundle-frags 10000000) [15].

### Gene Fusions

Gene fusions were detected from RNA and DNA using Integrate v0.2.0 [16], with default parameters.

### Data Deposition

All sequencing data are deposited in dbGAP study number [submission in progress]

### Heterogeneity analysis

Using the high depth of coverage from combining exome and WGS datasets for these tumors, we characterized the heterogeneity of each tumor specimen and compared it to the others in the series. Here, copy number-neutral variants and their attendant VAFs were clustered in five dimensions using the sciClone algorithm v1.1 [17] (parameters: minimumDepth=300, maximumClusters=15), followed by phylogeny reconstruction with clonEvol (https://github.com/hdng/clonevol).

### Neoantigen predictions

Somatic mutations and RNAseq data from tumors were input into our pVAC-Seq pipeline [18]. WGS data from the normal blood was used to identify the patient’s HLA haplotypes for class I, using HLAminer [19]. MHC class I binding predictions were generated through pVAC-Seq using NetMHC v3.4, as well as 5 other algorithms from the Immune Epitope Database and Analysis resource (IEDB, iedb.org): netMHC, netmhccons, netmhcpan, pickpocket, smm, and smmpmbec. Predictions were retained if the median score had an ic50 < 500 and better binding of the mutant peptide than the wild type (fold-change > 1). Results were then filtered to require expression of the mutant allele (FPKM > 1 and at least one variant-supporting read in the RNA). These combined data sets were used to identify neoantigenic peptide sequences in all five tumor samples, as illustrated in Fig 3c.

### Pathology methods

All the resection specimens (original and recurrences) were handled as regular surgical neuropathology cases. While hematoxylin and eosin stain and Ki-67 immunostain were performed on all the specimens, glial fibrillary acidic protein and epithelial membrane antigen were limited to the initial and 2014 resections.

## Results

### Mutational analysis of the matched normal sample

In order to determine whether the patient possessed a germline predisposition to cancer, we analyzed the sequence data obtained from her leukocyte-derived DNA (normal) and identified 176 protein-altering constitutional variants that were rare in the population and fell into highly-damaging classes of mutations (frameshifts, nonsense, nonstep, or splice-site) A full list of these is available at dbGAP accession xxxxxx). Variants were recognized in several genes known to be important for immune function, including splice site SNPs in *RAG1, HLA-DRB1,* and *HLA-DRB5,* as well as a nonsense mutation in *HLA-DRB5.* Several cancer relevant genes were also observed: splice-site alterations in *DDX3X* [20] and *MAD2L2* (alias: *REV7*) [21][22][23][24], in-frame insertions in *MNX1 [25]* and *ZFHX3, [26].* Some with direct glioma relevance were also observed: *FOXD1* (in frame deletion) [27][28][29], *BCL2L2* (SNP) [30], and *RYK (*frameshift insertion)[31]. The only predisposition-relevant gene was *MNX1 (*also known as *HLXB9)* which functions as an oncogene to promote pancreatic islet cell tumors in Multiple Endocrine Neoplasia Type 1 (MEN1) [32].

### Mutational landscape during disease progression

We identified 1,332 somatic mutations across the five resection specimens, 162 of which were in proteincoding regions, and 110 of which were non-silent (**Fig 2, S2 Table**). The primary tumor sample contained only one overtly cancer-related gene mutation, an expressed frameshift insertion in *MEN1* (K237fs) (**Table 1**). We also observed several large copy number alterations (CNAs) in this sample, including deletions of 6p, 15q, 22, and the first 22Mb of chromosome 1, that were shared with the recurrent tumors. (**S1 Figure, S3 Table)** Chromosome 11 was heavily rearranged, with multiple distinct regions of amplification and deletion, one of which deleted the second copy of *MEN1.* Integrated analysis of the DNA and RNA only detected gene fusion events in the fourth recurrence, likely due to significantly higher depth and quality of the RNA sequencing data from that sample (**S4 Table**). Many putative structural variants were detected in all samples (**S5 Table**).

**Fig 2.**
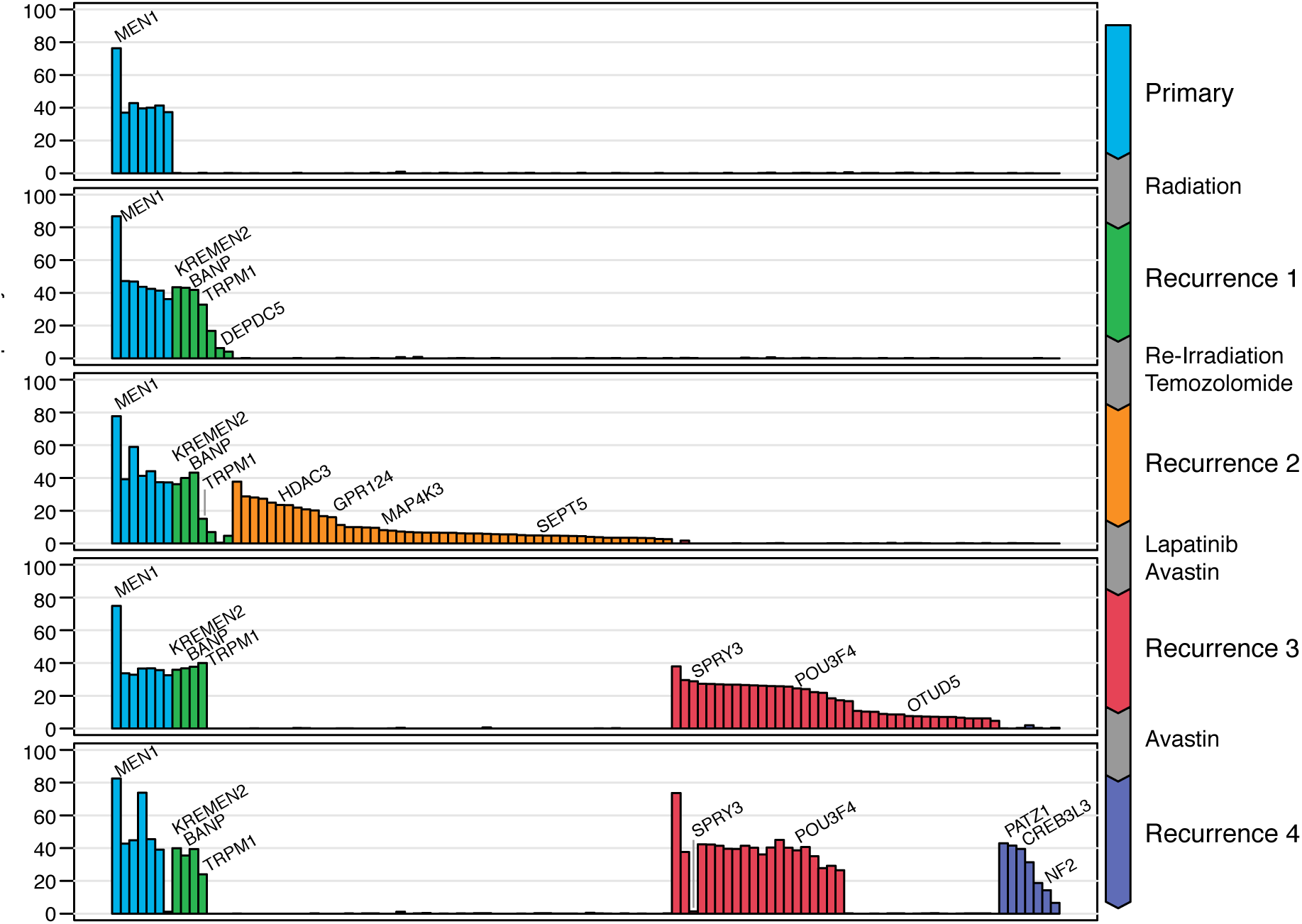
Variant allele fractions of non-silent mutations in protein-coding genes in all five resections.

**Table 1:** Functions and Cancer Relatedness of Mutated Genes

| Pathways/Gene | Function | References |
| --- | --- | --- |
| **Epigenetics** |  |  |
| *MEN1* | H3K4 trimethylation, DNA methylation | 23850066 |
| *GON4L* | HDAC1 interactor | 21454521 |
| *HDAC3* | Histone Deacetylase | 25313724 |
| *SETD9* | Histone methyltransferase | 20930037 |
| *BANP/SMAR1* | Recruits HDAC1 and deacetylation of H3K9 | 16166625 |
| *SUV39H1* | H3K9 trimethylase | 26807716 |
| *FAM208A* | Component of HUSH complex. Required for H3K9me3 | 26022416 |
| *POU3F4* | Epigenetics of neuronal fate | 23933087 |
| *PATZ1* | Regulates chromatin openness and pluripotency | 25515777 |
| *KAT6B* | Histone acetyltransferase | 26208904 |
| **Intracellular Signaling** |  |  |
| *KREMEN2* | Inhibitor of WNT signang | 20846389 |
| *DEPDC5* | Inhibitor of mTORC signaling | 23723238 |
| *NENF* | Neurotropic factor | 26042224 |
| *NET1* | rhoGEF required for proliferation | 23864709 |
| *SPRY3* | Inhibitor of FGF signaling | 18219583 |
| *VBP1* | VHL protein interactor | 23964080 |
| *ARHGAP32* | Inhibitor of RHOA, CDC42 and RAC1 signaling | 12857875 |
| *OTUD5* | Activator of p53 | 24143256 |
| *KLHL21* | Regulator of mitosis | 19995937 |
| *NF2* | Tumor suppressor, regulator of Hippo pathway | 25893302 |
| *LATS1* | Mediator of Hippo pathway, regulated by *NF2* | 25026211 |
| *MAP4K3* | Component of MAPK pathway regulator of LATS1 | 26437443 |
| *TRAF3* | TNFR-associated regulator of MAPK and NFκB pathways | 28098136 |
| **Metabolism** |  |  |
| *LETM1* | Mitochondrial structural protein | 25077561 |
| *TXNRD2* | Thioredoxin Reductase 2 | 25647640 |
| *HSD3B2* | Required for steroid biosynthesis | 22262841 |
| *BGAL* | Beta galactosidase | 23011886 |
| *MT-ND4* | Core component of mitochondrial NADH dehydrogenase | 25909222 |
| **Neuro-developmental disorders** |  |  |
| *SYNE1* | SYNE1 Ataxia | 27086870 |
| *GPR124* | CNS angiogenesis | 21071672 |
| *ARFGEF2* | Periventricular heterotopia | 23755938 |
| *CAGNG2* | Bipolar disorder | 25730879 |
| *SH3TC2* | Charcot Marie Tooth | 20028792 |
| **Other cancer related genes** |  |  |
| *NRBF2* | Breast cancer | 26073781 |
| *SLC39A11* | Ovarian cancer | 26091520 |
| *TRPM1* | Melanoma | 22897572 |

All SNVs, indels and CNAs found in the initial resection were retained in the first recurrence, which was diagnosed after radiation therapy and a 44 month interval from the first surgery. An additional 12 new protein-coding somatic mutations were identified in the recurrent tumor, including a nonsense mutation in *DEPDC5,* an inhibitor of mTORC signaling. Missense mutations were observed in *KREMEN2* (G165V), a gene that has been linked to melanoma, and in *BANP* (N223S), an epigenetic regulator. None are obviously expressed in this tumor, but the variants in both *KREMEN2* and *BANP* are expressed in subsequent tumors with higher quality and higher-depth RNAseq, so it is likely that these variants are expressed below our level of sensitivity in this resection sample. Mutated *DEPDC5* may have been undetectable due to undergoing nonsense-mediated decay.

The second recurrent tumor emerged after additional radiation therapy and treatment with temozolomide. It was resected and the genomic analysis of this specimen indicated that essentially all previously observed mutations were retained, with the exception of 2 low-VAF protein-coding variants from the previous recurrence, including loss of the *DEPDC5* nonsense mutation. An additional 66 protein-coding SNVs and indels were acquired, including a 19-bp frame-shift deletion in *GPR124* and low-VAF missense mutations in *SEPT5* (T260A), *MAP4K3* (F300S), and *KAT6B* (P1276L). Of these, only the *MAP43K* and *KAT6B* mutations were observably expressed. The copy number landscape was identical to the previous tumors, with the exception of a new homozygous deletion on chromosome 2p.

The third recurrence occurred after treatment with Avastin and Lapatinib. Genomic analysis of this resection specimen revealed that all coding mutations specific to the second recurrence, including the chromosome 2 copy number loss, were undetectable at the third recurrence. In contrast, virtually all mutations identified in first two resections persisted, the only exception being two low-VAF events in *MYH10* and *OR1L1.* 56 new protein-coding mutations were acquired, including missense mutations in *POU3F4* (P568T), an epigenetic regulator, *OTUD5* (P338L), a p53 activator, and *SPRY3* (R19C), a regulator of FGF signaling. None have been previously implicated in epenydymoma and their relevance for disease progression and therapy resistance is unclear.

The fourth recurrence was resected after continued Avastin treatment. In this sample, 29 of the proteincoding mutations newly acquired in the prior (third) recurrence were no longer detected, but 18 new proteincoding mutations were identified. These included nonsense mutations in *CREB3L3* and *NF2,* a gene previously linked to ependymoma. A missense mutation in the chromatin/transcriptional regulator *PATZ1* was also observed. In addition to the *NF2* mutation, we identified two point mutations that potentially impact Hippo pathway signaling in *LATS1* and *MAP4K3 [33][34].*

### Clonal heterogeneity and tumor evolution

To characterize the changing clonal architecture of this tumor, the variant allele fractions of copy-number neutral SNVs were clustered in 5 dimensions using the sciClone algorithm (**Fig 3A**). Eight clusters were detected, and the mutation spectrum for each was identified. The first recurrent tumor after radiation therapy was dominated by Cluster 2, which emerged from a population of cells undetectable in our analysis of the original biopsy data (with a sensitivity of about ~2% VAF). The mutation spectrum shows a notable decrease in C>T transitions in cluster 2, as compared to those in cluster 1 from the original tumor (**Fig 3B**). We attempted mutational signature analysis [35], but there were too few mutations to obtain statistically meaningful results.

**Fig 3:**
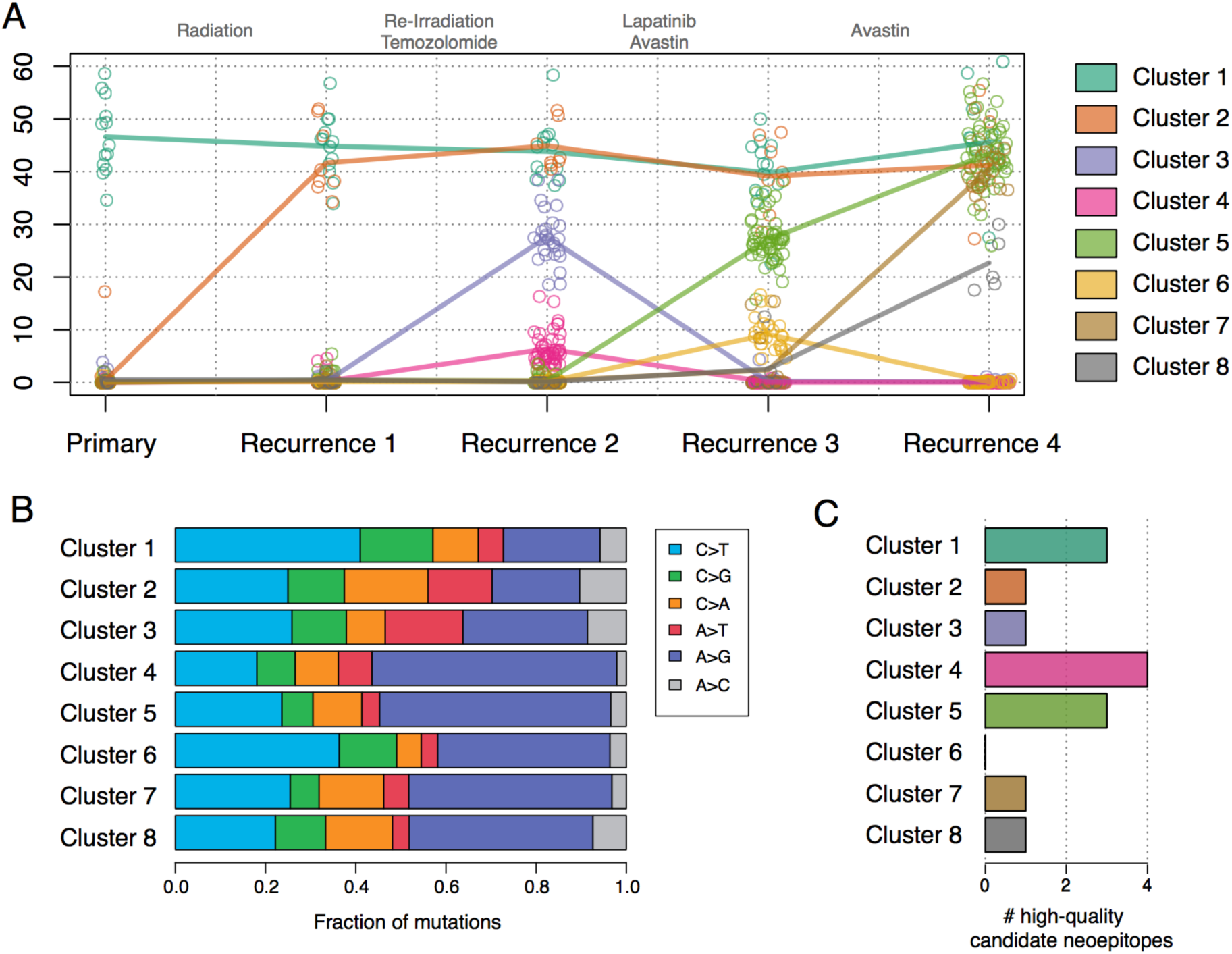
A) Subclonal clustering of the five tumor samples. Points represent the VAFs of individual SNVs at each timepoint, and lines connect the mean VAF of each cluster in each sample. B) mutation spectrum of each cluster C) the number of high-quality MHC Class I neoantigens found in each subclonal population.

In the second recurrence, following additional radiation and treatment with temozolomide, we identified the emergence of two new subclonal populations (clusters 3 and 4) that were likewise undetectable in the prior two samples. Cluster 4, and all subsequently appearing clusters, each have a significantly higher proportion of A>G transitions than the founding clone, a pattern consistent with temozolomide-induced mutagenesis (all p < 0.03, **S6 Table**) [36]. In the third recurrence, following Avastin treatment, both clusters 3 and 4 were undetectable, but clusters 5 and 6 emerged. Although Cluster 6 was cleared in the final resection sample we studied, Cluster 5 persisted and two rare subclonal populations expanded into clusters 7 and 8, which make up a substantial portion of the final tumor. Some mutations in these two clusters were just above the level of detection in Sample 4.

In addition to identifying mutations correlated with specific subclonal expansions, we also examined the expression of O6-methylguanine DNA methyltransferase *(MGMT),* which is known to drive brain tumor recurrence through increased expression in post-temozolomide lesions [37,38]. In this case, *MGMT* RNA expression levels were not clearly correlated with the emergence of post-temozolomide recurrences, suggesting that they relied upon alternative mechanisms of resistance (**S2 Figure**)

### Evolving landscape of targets for immunotherapy

In order to understand how the immunogenicity of this tumor evolved over the course of treatment, we applied the pVACSeq neoantigen prediction pipeline [18] to the protein-altering mutations that we observed in each tumor studied. The patient’s HLA haplotypes were inferred to be A*24:02, A*26:01, B*40:02, B*38:01, C*12:03, and C*03:05. We identified only 14 expressed mutations that produce “high-quality” predicted MHC Class I neoantigens (Fig 2C), which we define as having median binding affinity (ic50) of less than 500nM, and with a higher binding affinity to the mutant than the wildtype peptide (**S7 Table**). As overall mutation burden is highly correlated with neoantigen load, this is perhaps unsurprising. Only 3 neoantigens were present in the founding clone, while 11 of the 14 were specific to a subclonal population and therefore not present in all cells of the tumor.

## Discussion

These data reveal new insights into molecular features of ependymoma and the effects of therapy on tumor evolution. Most notable is our identification of *MEN1* mutation as a putative driver mutation, of potential mechanisms that contributed to tumor evolution, and of an observed *GPR124* mutation that may serve as a prognostic indicator of response to Avastin.

### MEN1 mutation and copy number loss

*MEN1* mutations occur infrequently in ependymoma, both in the context of familial MEN 1 syndrome and sporadically [39][40][41][42]. Whereas the genetics of supratentorial and infratentorial ependymoma, as well as the genetics of low grade (WHO grades I and II) and high grade (WHO grade III) ependymoma are most often reported to be distinct, MEN1 mutations occur, albeit rarely, in ependymomas of any grade and location. As is the case with the pathognomonic neoplasms of MEN1 syndrome, there is biallelic disruption of MEN1 in ependymomas. Two prior reports of recurrent ependymomas suggested that MEN1 mutation was associated with recurrence and progression from grade II to III [43][41]. Ours is the first report to define the clonal architecture of primary and recurrent ependymoma and to identify MEN1 mutation as a feature of the founding clone.

It is important to consider the potential interaction between the altered *MEN1* protein and germline altered cancer relevant proteins in the genesis of the initial tumor clone. In this regard, *MNX1* expression is known to be regulated by Menin, the protein product of *MEN1.* Though it has been reported that loss of Menin function increases *MNX1* expression [32]], and *MNX1* has also been shown to function as an oncogene in driving pancreatic islet cell tumorigenesis [44][45][46], we did not observe substantial expression of *MNX1* in any of our tumors.

### MEN1 dependent mechanisms in tumor evolution

Biochemical and genetic studies of Menin, the protein product of *MEN1,* have revealed complex and cell type specific functions that include roles in epigenetics, transcriptional regulation and regulation of intracellular signaling [47]. Of particular relevance to known ependymoma mechanisms are Menin’s roles in DNA methylation and intracellular signaling pathways known to be involved in glioma biology, described below.

In both human *MEN1* syndrome-associated tumors and mouse models of *MEN1,* loss of menin function results in increased DNA methyltransferase 1 (DNMT1) activity and global increases in CpG island methylation [48]. This CpG Island Methylator Phenotype (CIMP) is a known characteristic feature of colon cancer and a subset of high-grade gliomas, as well as a subset of ependymoma [49][5]. Tumor suppressors known to be silenced by menin loss and to be relevant to glioma biology include *CDKN1B, CDKN2A, APC and RASSF1A [50][51]* (PMID:16195383). The importance of altered epigenetic mechanisms to the biology of the ependymoma recurrences studied here is also suggested by the accumulation of mutations in additional epigenetic regulators through the course of its evolution *(POU3F4, GON4L, HDAC3, SETD9, SUV39H1, BANP/SMAR1, FAM208A, PATZ1, KAT6B*).

Loss of menin function also is known to directly regulate multiple intracellular signaling pathways with established roles in gliomagenesis. Most relevant to glioma biology are the enhancing effects of menin loss on the activation of the RAS [52], MAP kinase [53], PI3 kinase [54], Sonic Hedgehog [55], Wnt [56] and TGF-b [57] signaling pathways. The importance of some of these mechanisms to the biology of this ependymoma is also suggested by the accumulation of mutations in additional regulators of their activation: MAPK *(SPRY3* [58]), PI3K (DEPDC5 [58,59], WNT *(KREMEN2* [60][61], *NET1 [62], GPR124 [63]).*

In addition to the glioma-relevant pathways regulated by Menin, the Hippo pathway stands out as a compelling potential driver of the fourth recurrence, induced by new mutations in *LATS1* (missense) and in *NF2* (nonsense). *NF2* loss-of-function mutations have been previously associated with spinal ependymomas [64], and recent evidence suggests that *YAP1,* the nuclear target of Hippo signaling, mediates aberrant proliferation upon *NF2* loss during tumorigenesis [65]. Furthermore, oncogenic *YAP1* activation occurs as a consequence of a loss in NF2-dependent inactivation of *LATS1,* a key inhibitor of *YAP1* [33,65]. Decreased *LATS1* activity has also been associated with glioma progression [33,65,66].

### *GPR124* mutation as a potential prognostic indicator of Avastin Response

Additional striking features in the evolution of this ependymoma include the persistence of the founding clone and the unique response of the subclone arising in Recurrence 4 to Avastin and Lapatinib. This is the only substantial subclone to be completely eradicated by any element of treatment, suggesting an important relationship between this therapy and a target in this clone. Among the compelling target mediators of response or biomarkers of response is the mutation in *GPR124.* This orphan member of the adhesion G protein coupled receptor family is required specifically for the development of the brain vasculature [67][68][69] in a VEGF-dependent manner, and GPR124 may be a biomarker of Avastin response [70]. GPR124 activates canonical Wnt signaling which, as described above, is normally directly inhibited by MEN 1 and KREMEN2. The genes for both of these proteins were mutated in this tumor, suggesting a model for avastin response that might involve enhanced activation of a VEGF-Wnt axis. Additional study of samples before and after Avastin treatment is important to explore this possibility further.

### Changes to the neoantigen landscape during progression

Overall, the number of expressed MHC Class I neoantigens that we predicted was low, as expected in a tumor with only 110 protein-altering mutations. The presence of a relatively high burden in the founding clone suggests that mechanisms of immune evasion were already present when the initial tumor presented, and may explain why there was no relationship between neoantigen load and subclonal response in subsequent tumors. This is supported by the observation that expression markers of T-cell activation were low in all five tumors.

### Future Directions

These results suggest that radiation and chemotherapy contributed to the increasing complexity of this tumor by both adding to the mutational burden and expanding the subclonal architecture. Determining whether this natural history is generally true in ependymoma progression, and what impact therapy-induced tumor evolution has on outcome, is an important area of investigation with the potential to alter how we treat patients with completely resected supratentorial ependymoma. In the largest published study of ependymoma outcome involving 282 patients, gross total resection (GTR) was the only prognostic factor associated with increased survival [71]. In this analysis, GTR and post-surgical radiation therapy were associated with a shorter progression-free survival than GTR alone.

These clinical observations together with the sequencing-based characterizations presented here suggest that under some circumstances, adjuvant therapy may not be providing a benefit, and indeed may hasten recurrence by promoting molecular diversification of the tumor. We propose that this phenomenon be studied prospectively by sequencing completely resected supratentorial ependymomas and subsequent relapses to determine (a) the circumstances under which radiation therapy eradicates the founding clone and results in cure and (b) the circumstances under which tumors are resistant and instead exhibit an increase in their genomic complexity. Ultimately, it may be prudent to initially observe those patients with complete resections without additional therapy or to treat those patients whose tumors are likely to evolve in response to radiation therapy with targeted agents only as dictated by genomic analysis.

Finally, it will be important to investigate further the utility of genomic characterization to inform therapeutic options in this disease type. Although not all of the variants we identified were “druggable” in the classical sense, a subset were found to comprise predicted high affinity neoantigen targets that, ultimately, formed the basis of a polyvalent personalized vaccine, administered after recurrence 4. Although the efficacy of these treatments awaits large scale studies that are ongoing, our case highlights the potential to consider the pursuit of a personalized vaccine in extremely challenging settings of multiple recurrent disease such as the one herein, where few to no other options exist.

## Acknowledgements

The authors would like to acknowledge early analysis efforts on this project from Charles Lu and assistance with INTEGRATE from Jin Zhang. They also gratefully acknowledge the sample intake, project tracking and data production teams at the McDonnell Genome Institute. Funding for this project was generously provided from the McDonnell Genome Institute endowment and from the Robert E. and Louise F. Dunn Distinguished Professorship endowment to Washington University School of Medicine. We gratefully acknowledge the patient and her family.

---

**S1 Figure:** Copy number alterations found in all 5 samples.

**S2 Figure:** *MGMT* expression from RNA sequencing (units are FPKM). The 4th recurrence sample is presented for completeness, but was sequenced after exome capture, (see S1 Table), which prevents meaningful comparisons to the other samples.

**S1 Table:** DNA and RNA sequencing coverage

**S2 Table:** Somatic variants and readcounts from both DNA and RNA in all resections.

**S3 Table:** Copy number alterations and regions of loss-of-heterozygosity identified in all 5 resections.

**S4 Table:** Structural variants identified in all 5 resections

**S6 Table:** Fraction of A>G transitions in each subclone. P-value calculated by comparing the proportion of A>G mutations in each cluster to those in the founding clone (Cluster 1) using Fisher’s exact test. Multiple-testing correction was applied using the Benjamini-Hochberg method as implemented in the p.adjust() function in R.

**S7 Table:** Predicted high-quality neoantigens identified in all 5 resections.

## References

1. Khatua S, Ramaswamy V, Bouffet E. Current therapy and the evolving molecular landscape of paediatric ependymoma. Eur J Cancer. 2017;70: 34–41.

2. Dorfer C, Tonn J, Rutka JT. Ependymoma: a heterogeneous tumor of uncertain origin and limited therapeutic options. Handb Clin Neurol. 2016; 134: 417–431.

3. Pajtler KW, Witt H, Sill M, Jones DTW, Hovestadt V, Kratochwil F, et al. Molecular Classification of Ependymal Tumors across All CNS Compartments, Histopathological Grades, and Age Groups. Cancer Cell. 2015;27: 728–743.

4. Pajtler KW, Mack SC, Ramaswamy V, Smith CA, Witt H, Smith A, et al. The current consensus on the clinical management of intracranial ependymoma and its distinct molecular variants. Acta Neuropathol. 2017; 133: 5–12.

5. Mack SC, Witt H, Piro RM, Gu L, Zuyderduyn S, Stütz AM, et al. Epigenomic alterations define lethal CIMP-positive ependymomas of infancy. Nature. 2014;506: 445–450.

6. Griffith M, Griffith OL, Smith SM, Ramu A, Callaway MB, Brummett AM, et al. Genome Modeling System: A Knowledge Management Platform for Genomics. PLoS Comput Biol. 2015; 11: e1004274.

7. Li H, Handsaker B, Wysoker A, Fennell T, Ruan J, Homer N, et al. The Sequence Alignment/Map format and SAMtools. Bioinformatics. 2009;25: 2078–2079.

8. Larson DE, Harris CC, Chen K, Koboldt DC, Abbott TE, Dooling DJ, et al. SomaticSniper: identification of somatic point mutations in whole genome sequencing data. Bioinformatics. 2012;28: 311–317.

9. Koboldt DC, Zhang Q, Larson DE, Shen D, McLellan MD, Lin L, et al. VarScan 2: somatic mutation and copy number alteration discovery in cancer by exome sequencing. Genome Res. 2012;22: 568–576.

10. Saunders CT, Wong WSW, Swamy S, Becq J, Murray LJ, Cheetham RK. Strelka: accurate somatic small-variant calling from sequenced tumor-normal sample pairs. Bioinformatics. 2012;28: 1811–1817.

11. Cibulskis K, Lawrence MS, Carter SL, Sivachenko A, Jaffe D, Sougnez C, et al. Sensitive detection of somatic point mutations in impure and heterogeneous cancer samples. Nat Biotechnol. 2013;31: 213–219.

12. McKenna A, Hanna M, Banks E, Sivachenko A, Cibulskis K, Kernytsky A, et al. The Genome Analysis Toolkit: a MapReduce framework for analyzing next-generation DNA sequencing data. Genome Res. 2010;20: 1297–1303.

13. Ye K, Schulz MH, Long Q, Apweiler R, Ning Z. Pindel: a pattern growth approach to detect break points of large deletions and medium sized insertions from paired-end short reads. Bioinformatics. 2009;25: 2865–2871.

14. Chen K, Wallis JW, McLellan MD, Larson DE, Kalicki JM, Pohl CS, et al. BreakDancer: an algorithm for high-resolution mapping of genomic structural variation. Nat Methods. 2009;6: 677–681.

15. Trapnell C, Roberts A, Goff L, Pertea G, Kim D, Kelley DR, et al. Differential gene and transcript expression analysis of RNA-seq experiments with TopHat and Cufflinks. Nat Protoc. 2012;7: 562–578.

16. Zhang J, White NM, Schmidt HK, Fulton RS, Tomlinson C, Warren WC, et al. INTEGRATE: gene fusion discovery using whole genome and transcriptome data. Genome Res. 2016;26: 108–118.

17. Miller CA, White BS, Dees ND, Griffith M, Welch JS, Griffith OL, et al. SciClone: inferring clonal architecture and tracking the spatial and temporal patterns of tumor evolution. PLoS Comput Biol. 2014;10: e1003665.

18. Hundal J, Carreno BM, Petti AA, Linette GP, Griffith OL, Mardis ER, et al. pVAC-Seq: A genome-guided in silico approach to identifying tumor neoantigens. Genome Med. 2016;8: 11.

19. Warren RL, Choe G, Freeman DJ, Castellarin M, Munro S, Moore R, et al. Derivation of HLA types from shotgun sequence datasets. Genome Med. 2012;4: 95.

20. Dahlin AM, Hollegaard MV, Wibom C, Andersson U, Hougaard DM, Deltour I, et al. CCND2, CTNNB1, DDX3X, GLI2, SMARCA4, MYC, MYCN, PTCH1, TP53, and MLL2 gene variants and risk of childhood medulloblastoma. J Neurooncol. 2015; 125: 75–78.

21. Okina S, Yanagisawa N, Yokoyama M, Sakurai Y, Numata Y, Umezawa A, et al. High expression of REV7 is an independent prognostic indicator in patients with diffuse large B-cell lymphoma treated with rituximab. Int J Hematol. 2015;102: 662–669.

22. Xu G, Chapman JR, Brandsma I, Yuan J, Mistrik M, Bouwman P, et al. REV7 counteracts DNA double-strand break resection and affects PARP inhibition. Nature. 2015;521: 541–544.

23. Niimi K, Murakumo Y, Watanabe N, Kato T, Mii S, Enomoto A, et al. Suppression of REV7 enhances cisplatin sensitivity in ovarian clear cell carcinoma cells. Cancer Sci. 2014;105: 545–552.

24. Boersma V, Moatti N, Segura-Bayona S, Peuscher MH, van der Torre J, Wevers BA, et al. MAD2L2 controls DNA repair at telomeres and DNA breaks by inhibiting 5’ end resection. Nature. 2015;521: 537–540.

25. Das M. MNX1: a novel prostate cancer oncogene. Lancet Oncol. 2016; 17: e521.

26. Mabuchi M, Kataoka H, Miura Y, Kim T-S, Kawaguchi M, Ebi M, et al. Tumor suppressor, AT motif binding factor 1 (ATBF1), translocates to the nucleus with runt domain transcription factor 3 (RUNX3) in response to TGF-beta signal transduction. Biochem Biophys Res Commun. 2010;398: 321–325.

27. Cheng P, Wang J, Waghmare I, Sartini S, Coviello V, Zhang Z, et al. FOXD1-ALDH1A3 Signaling Is a Determinant for the Self-Renewal and Tumorigenicity of Mesenchymal Glioma Stem Cells. Cancer Res. 2016;76: 7219–7230.

28. Koga M, Matsuda M, Kawamura T, Sogo T, Shigeno A, Nishida E, et al. Foxd1 is a mediator and indicator of the cell reprogramming process. Nat Commun. 2014;5: 3197.

29. Gao Y-F, Zhu T, Mao X-Y, Mao C-X, Li L, Yin J-Y, et al. Silencing of Forkhead box D1 inhibits proliferation and migration in glioma cells. Oncol Rep. 2017;37: 1196–1202.

30. Chung HJ, Choi YE, Kim ES, Han Y-H, Park M-J, Bae IH. miR-29b attenuates tumorigenicity and stemness maintenance in human glioblastoma multiforme by directly targeting BCL2L2. Oncotarget. 2015;6: 18429–18444.

31. Adamo A, Fiore D, De Martino F, Roscigno G, Affinito A, Donnarumma E, et al. RYK promotes the stemness of glioblastoma cells via the WNT/ β-catenin pathway. Oncotarget. 2017; doi: 10.18632/oncotarget.14564

32. Scacheri PC, Davis S, Odom DT, Crawford GE, Perkins S, Halawi MJ, et al. Genome-wide analysis of menin binding provides insights into MEN1 tumorigenesis. PLoS Genet. 2006;2: e51.

33. Oh J-E, Ohta T, Satomi K, Foll M, Durand G, McKay J, et al. Alterations in the NF2/LATS1/LATS2/YAP Pathway in Schwannomas. J Neuropathol Exp Neurol. 2015;74: 952–959.

34. Meng Z, Moroishi T, Mottier-Pavie V, Plouffe SW, Hansen CG, Hong AW, et al. MAP4K family kinases act in parallel to MST1/2 to activate LATS1/2 in the Hippo pathway. Nat Commun. 2015;6: 8357.

35. Rosenthal R, McGranahan N, Herrero J, Taylor BS, Swanton C. DeconstructSigs: delineating mutational processes in single tumors distinguishes DNA repair deficiencies and patterns of carcinoma evolution. Genome Biol. 2016; 17: 31.

36. Bodell WJ, Gaikwad NW, Miller D, Berger MS. Formation of DNA adducts and induction of lacI mutations in Big Blue Rat-2 cells treated with temozolomide: implications for the treatment of low-grade adult and pediatric brain tumors. Cancer Epidemiol Biomarkers Prev. 2003;12: 545–551.

37. Bocangel DB, Finkelstein S, Schold SC, Bhakat KK, Mitra S, Kokkinakis DM. Multifaceted resistance of gliomas to temozolomide. Clin Cancer Res. 2002;8: 2725–2734.

38. Hegi ME, Diserens A-C, Gorlia T, Hamou M-F, de Tribolet N, Weller M, et al. MGMT gene silencing and benefit from temozolomide in glioblastoma. N Engl J Med. 2005;352: 997–1003.

39. Lee C-H, Chung CK, Ohn JH, Kim CH. The Similarities and Differences between Intracranial and Spinal Ependymomas: A Review from a Genetic Research Perspective. J Korean Neurosurg Soc. 2016;59: 83–90.

40. Al-Salameh A, François P, Giraud S, Calender A, Bergemer-Fouquet A-M, de Calan L, et al. Intracranial ependymoma associated with multiple endocrine neoplasia type 1. J Endocrinol Invest. 2010;33: 353–356.

41. Urioste M, Martínez-Ramírez A, Cigudosa JC, Colmenero I, Madero L, Robledo M, et al. Complex cytogenetic abnormalities including telomeric associations and MEN1 mutation in a pediatric ependymoma. Cancer Genet Cytogenet. 2002;138: 107–110.

42. Kato H, Uchimura I, Morohoshi M, Fujisawa K, Kobayashi Y, Numano F, et al. Multiple endocrine neoplasia type 1 associated with spinal ependymoma. Intern Med. 1996;35: 285–289.

43. Funayama T, Sakane M, Yoshizawa T, Takeuchi Y, Ochiai N. Tanycytic ependymoma of the filum terminale associated with multiple endocrine neoplasia type 1: first reported case. Spine J. 2013;13: e49–54.

44. Desai SS, Kharade SS, Parekh VI, Iyer S, Agarwal SK. Pro-oncogenic Roles of HLXB9 Protein in Insulinoma Cells through Interaction with Nono Protein and Down-regulation of the c-Met Inhibitor Cblb (Casitas B-lineage Lymphoma b). J Biol Chem. 2015;290: 25595–25608.

45. Desai SS, Modali SD, Parekh VI, Kebebew E, Agarwal SK. GSK-3β protein phosphorylates and stabilizes HLXB9 protein in insulinoma cells to form a targetable mechanism of controlling insulinoma cell proliferation. J Biol Chem. 2014;289: 5386–5398.

46. Shi K, Parekh VI, Roy S, Desai SS, Agarwal SK. The embryonic transcription factor Hlxb9 is a menin interacting partner that controls pancreatic β-cell proliferation and the expression of insulin regulators. Endocr Relat Cancer. 2013;20: 111–122.

47. Matkar S, Thiel A, Hua X. Menin: a scaffold protein that controls gene expression and cell signaling. Trends Biochem Sci. 2013;38: 394–402.

48. Yuan Z, Sánchez Claros C, Suzuki M, Maggi EC, Kaner JD, Kinstlinger N, et al. Loss of MEN1 activates DNMT1 implicating DNA hypermethylation as a driver of MEN1 tumorigenesis. Oncotarget. 2016;7: 12633–12650.

49. Rogers HA, Kilday J-P, Mayne C, Ward J, Adamowicz-Brice M, Schwalbe EC, et al. Supratentorial and spinal pediatric ependymomas display a hypermethylated phenotype which includes the loss of tumor suppressor genes involved in the control of cell growth and death. Acta Neuropathol. 2012; 123: 711–725.

50. Lindberg D, Akerström G, Westin G. Evaluation of CDKN2C/p18, CDKN1B/p27 and CDKN2B/p15 mRNA expression, and CpG methylation status in sporadic and MEN 1-associated pancreatic endocrine tumours. Clin Endocrinol. 2008;68: 271–277.

51. Juhlin CC, Kiss NB, Villablanca A, Haglund F, Nordenström J, Höög A, et al. Frequent promoter hypermethylation of the APC and RASSF1A tumour suppressors in parathyroid tumours. PLoS One. 2010;5: e9472.

52. Wu Y, Feng Z-J, Gao S-B, Matkar S, Xu B, Duan H-B, et al. Interplay between menin and K-Ras in regulating lung adenocarcinoma. J Biol Chem. 2012;287: 40003–40011.

53. Gallo A, Cuozzo C, Esposito I, Maggiolini M, Bonofiglio D, Vivacqua A, et al. Menin uncouples Elk-1, JunD and c-Jun phosphorylation from MAP kinase activation. Oncogene. 2002;21: 6434–6445.

54. Wang Y, Ozawa A, Zaman S, Prasad NB, Chandrasekharappa SC, Agarwal SK, et al. The tumor suppressor protein menin inhibits AKT activation by regulating its cellular localization. Cancer Res. 2011;71: 371–382.

55. Gurung B, Feng Z, Iwamoto DV, Thiel A, Jin G, Fan C-M, et al. Menin epigenetically represses Hedgehog signaling in MEN1 tumor syndrome. Cancer Res. 2013;73: 2650–2658.

56. Cao Y, Liu R, Jiang X, Lu J, Jiang J, Zhang C, et al. Nuclear-cytoplasmic shuttling of menin regulates nuclear translocation of {beta}-catenin. Mol Cell Biol. 2009;29: 5477–5487.

57. Mould AW, Duncan R, Serewko-Auret M, Loffler KA, Biondi C, Gartside M, et al. Global expression profiling of sex cord stromal tumors from Men1 heterozygous mice identifies altered TGF-beta signaling, decreased Gata6 and increased Csf1r expression. Int J Cancer. 2009;124: 1122–1132.

58. Cabrita MA, Christofori G. Sprouty proteins, masterminds of receptor tyrosine kinase signaling. Angiogenesis. 2008; 11: 53–62.

59. Bar-Peled L, Chantranupong L, Cherniack AD, Chen WW, Ottina KA, Grabiner BC, et al. A Tumor suppressor complex with GAP activity for the Rag GTPases that signal amino acid sufficiency to mTORC1. Science. 2013;340: 1100–1106.

60. Orlow S, Yasunami R, Boitard C, Bach JF. [Early induction of diabetes in NOD mice by streptozotocin]. C R Acad Sci III. 1987;304: 77–78.

61. Mao B, Wu W, Davidson G, Marhold J, Li M, Mechler BM, et al. Kremen proteins are Dickkopf receptors that regulate Wnt/beta-catenin signalling. Nature. 2002;417: 664–667.

62. Wei S, Dai M, Liu Z, Ma Y, Shang H, Cao Y, et al. The guanine nucleotide exchange factor Net1 facilitates the specification of dorsal cell fates in zebrafish embryos by promoting maternal β-catenin activation. Cell Res. 2017;27: 202–225.

63. Posokhova E, Shukla A, Seaman S, Volate S, Hilton MB, Wu B, et al. GPR124 functions as a WNT7-specific coactivator of canonical β-catenin signaling. Cell Rep. 2015;10: 123–130.

64. Lee C-H, Chung CK, Kim CH. Genetic differences on intracranial versus spinal cord ependymal tumors: a meta-analysis of genetic researches. Eur Spine J. 2016;25: 3942–3951.

65. Shi Y, Bollam SR, White SM, Laughlin SZ, Graham GT, Wadhwa M, et al. Rac1-Mediated DNA Damage and Inflammation Promote Nf2 Tumorigenesis but Also Limit Cell-Cycle Progression. Dev Cell. 2016;39: 452–465.

66. Ji T, Liu D, Shao W, Yang W, Wu H, Bian X. Decreased expression of LATS1 is correlated with the progression and prognosis of glioma. J Exp Clin Cancer Res. 2012;31: 67.

67. Kuhnert F, Mancuso MR, Shamloo A, Wang H-T, Choksi V, Florek M, et al. Essential regulation of CNS angiogenesis by the orphan G protein-coupled receptor GPR124. Science. 2010;330: 985–989.

68. Cullen M, Elzarrad MK, Seaman S, Zudaire E, Stevens J, Yang MY, et al. GPR124, an orphan G protein-coupled receptor, is required for CNS-specific vascularization and establishment of the blood-brain barrier. Proc Natl Acad Sci U S A. 2011;108: 5759–5764.

69. Zhou Y, Nathans J. Gpr124 controls CNS angiogenesis and blood-brain barrier integrity by promoting ligand-specific canonical wnt signaling. Dev Cell. 2014;31: 248–256.

70. Wang Y, Cho S-G, Wu X, Siwko S, Liu M. G-protein coupled receptor 124 (GPR124) in endothelial cells regulates vascular endothelial growth factor (VEGF)-induced tumor angiogenesis. Curr Mol Med. 2014;14: 543–554.

71. Vera-Bolanos E, Aldape K, Yuan Y, Wu J, Wani K, Necesito-Reyes MJ, et al. Clinical course and progression-free survival of adult intracranial and spinal ependymoma patients. Neuro Oncol. 2015;17: 440–447.

